# Genetic diversity and population structure of soybean (*Glycine max* (L.) Merril) germplasm

**DOI:** 10.1101/2024.10.02.616345

**Authors:** Tenena Silue, Paterne Angelot Agre, Bunmi Olasanmi, Adeyinka Saburi Adewumi, Idris Ishola Adejumobi, Abush Tesfaye Abebe

## Abstract

Soybean (*Glycine max* (L.) Merril) is a significant legume crop for oil and protein. However, its yield in Africa is less than half the global average resulting in low production, which is inadequate for satisfying the continent’s needs. To address this disparity in productivity, it is crucial to develop new high-yielding cultivars by utilizing the genetic diversity of existing germplasms. Consequently, the genetic diversity and population structure of various soybean accessions were evaluated in this study. In pursuit of this objective, a collection of 147 soybean accessions was genotyped via the Diversity Array Technology Sequencing method. This method enables high-throughput analysis of single-nucleotide polymorphisms (SNPs), resulting in the identification of 7,083 high-quality SNPs distributed throughout the soybean genome. The average values observed for polymorphism information content (PIC), minor allele frequency, expected heterozygosity and observed heterozygosity were 0.277, 0.254, 0.344, and 0.110, respectively. The soybean genotypes were categorized into four groups on the basis of model-based population structure, principal component analysis, and discriminant analysis of the principal component. Alternatively, hierarchical clustering was used to organize the accessions into three distinct clusters. Analysis of molecular variance indicated that the genetic variance within the populations exceeded the variance among them. The insights gained from this study will assist breeders in selecting parental lines for genetic recombination. Overall, this study provides valuable information regarding soybean genetic diversity and lays the groundwork for conservation and genetic enhancement initiatives.

## Introduction

Soybean (*Glycine max* (L.) Merril) is a self-pollinated crop from the Fabaceae family with a diploid chromosome number of 2n = 40 [1]. It is one of the world’s major legumes and oil crops in terms of production and trade [2]. Soybean contains approximately 38–42% high-quality protein and 18–20% oil rich in essential fatty acids [3]. In Nigeria, soybean is used to produce nutritious drinks known as “soymilk” and “awara” (soybean cake). It is also a crucial ingredient in poultry and fish feed and is also used in infant meals. The oil is utilized in cooking and as a base for mayonnaise, margarine, salad dressings, and shortening [4].

Globally, soybean cultivation covers approximately 121 million hectares, with an estimated total production of 334 million tons annually [5]. The top three producers (Brazil, the United States, and Argentina) together contribute 73% of the world’s production. In Africa, approximately 2.55 million hectares are dedicated to soybean cultivation, with an average productivity of 1,348 kg per hectare. South Africa, Nigeria, and Zambia are the leading producers on the continent, with annual production rates of 1.32, 0.73, and 0.35 million tons, respectively [6].

Soybean cultivation in Africa typically yields less than 1.5 t. ha^-1^, which is significantly below the potential yield of over 3 t. ha^-1^ [7]. This low productivity might be attributed to various factors, including the limited availability of high-yielding and climatically resilient improved varieties, poor soil fertility, diseases and pests, high pod shattering, inadequate agronomic practices, and particularly drought caused by inconsistent rainfall [8]. Therefore, there is a need for improved soybean varieties that are resilient to these biotic and abiotic stresses [9]. On the other hand, the widespread use of only a few elite cultivars by farmers has significantly contributed to the decline in the diversity of crop germplasm, which can contribute to resilience to adverse climatic factors amidst the increasing occurrence of extreme climatic conditions [5]. Furthermore, genetic diversity in many crops has decreased over time as commercial plant breeding focuses on enhancing one or a few traits and/or uses a limited number of exceptional genotypes to create a breeding population [10].

Exploiting and conserving crop genetic diversity is essential, whereas assessing genetic diversity within germplasms is vital for developing new cultivars with desirable traits [11]. Understanding a crop’s genetic diversity is necessary for expanding a core collection and enhancing germplasm utilization in breeding programs [2]. Moreover, identifying genes that regulate beneficial biological processes in existing populations (such as breeding lines, cultivars, landraces, and wild relatives) is crucial for improving current varieties and promoting sustainable farming practices [12]. Additionally, by understanding genetic variability within and between plant populations, breeders can better understand genetic exchange or gene flow within populations [13].

Crop variability can be assessed at both the phenotypic and genotypic levels via the use of statistical methods to separate phenotypic or genetic descriptors into genetic or environmental components [14]. While morphological markers detect diversity among genotypes, they are less effective than DNA markers because of their subjective nature, limited number and, susceptibility to environmental factors [9]. The use of these markers to evaluate genetic diversity aids in the efficient utilization of germplasms for conservation and crop yield enhancement. According to Fischer et al. [15], single polymorphism nucleotide (SNP) markers are among the most effective genetic markers for identifying variations in crop varieties. Recently, the adoption of Diversity Array Technology (DArT), an advanced genotyping by sequencing (GBS) platform, has significantly reduced the time and cost associated with generating high-density SNP information [14]. Consequently, diverse studies of numerous crop species have shown greater coverage and fewer missing data when DArT is used than when other GBS platforms are used [16].

The soybean breeding programme at the International Institute of Tropical Agriculture (IITA) in Ibadan, Nigeria, has conducted phenotypic assessments on 1800 soybean accessions, evaluating key traits such as disease resistance, pod-shattering tolerance, early and medium maturity, efficient natural nodulation, lodging tolerance, and yield improvement [2, 17]. Despite the potential benefits of using DArT-SNP technology, there has been limited application of this method in assessing the genetic diversity of the IITA soybean breeding program germplasm. The lack of comprehensive genetic data hampers the development of improved soybean varieties that are better adapted to local environmental conditions and resistant to pests and diseases. Furthermore, the absence of detailed population structure analysis limits our understanding of the genetic relationships and evolutionary history of the IITA soybean gene pool.

## Materials and methods

### Plant materials, planting, and leaf sampling

A total of 147 soybean accessions, comprised of 130 genotypes from the IITA soybean breeding program, and 17 genotypes sourced from the USDA genetic resource center (14 genotypes), Ghana (1 genotype), Uganda (1 genotype), and a private seed company (SeedCo) (1 genotype) (list of germplasms, S1 Table), were selected and utilized for a molecular-based diversity assessment.

The 147 soybean accessions were sown and grown to the seedling stage in a screen house at IITA station Ibadan, Nigeria at 243 m.a.s., 7°30′8″N longitude and 3°54′37″E latitude. Three weeks after planting, five-leaf discs 5 mm in diameter from young and healthy leaves were collected via a biopsy curette from the leaf blades of each of the 147 genotypes. The leaf samples were placed into 96-well collection plates (12 × 8-strip tubes per 96-deep well plate). The freeze-dried leaf samples were sent to Diversity Array Technology (DArT)®, Canberra, Australia, for DNA extraction, library construction, and SNP marker development.

### SNP Marker Quality Control

Single-row format data from DArT were initially converted into HapMap and variant call format (VCF) formats using KDcompute (https://kdcompute.seqart.net/kdcompute, accessed on 07/06/2024). SNP-derived markers were then first filtered using PLINK 1.9 and VCFtools, on the basis of the call rate of the raw data [18]. The SNP markers with call rates ranging from 0.80 to 1.0 were selected for further quality control analyses. Duplicate SNP markers across the chromosomes were removed. This process involved removing markers with minor allele frequencies of less than 5%, markers and genotypes with more than 20% missing data, and those with a low coverage read depth of less than 5 [20,21].

### Statistical Analyses

The structure and pattern of genetic diversity within soybean genotypes were assessed via genotypic data generated on the basis of SNP markers. VCFtools and PLINK 1.9 were used to estimate summary statistics such as observed and expected heterozygosity, minor allele frequency (MAF), and polymorphic information content (PIC). The SNP distribution and density plot of the SNP markers across the 20 chromosomes of the soybean genome was constructed via the CMplot package [22]. The SNP markers data were subjected to population structure analysis following the method described by Agre et al. [23]. By testing cluster numbers ranging from 2 to 10, the optimal number of clusters was identified through k-means analysis, employing cross-validation on the basis of the Bayesian information criterion (BIC). Each soybean genotype was assigned to its respective cluster if it had at least 70% ancestry probability. Genotypes with less than 70% ancestry were considered as admixed. The diversity pattern revealed through population structure analysis was further supported by discriminant analysis of the principal component (DAPC) via the Adegenet package in R [24]. DAPC, which uses the k-means clustering method, aims to minimize variance within clusters while maximizing variance between clusters [25]. Pairwise genetic dissimilarity distances (identity-by-state, IBS) were calculated via the Jaccard method, implemented in the Philentropy R package [26]. A Ward’s minimum variance hierarchical cluster dendrogram was then constructed from the Jaccard dissimilarity matrix using the Analyses of Phylogenetics and Evolution (APE) package in R [27]. Principal component analysis (PCA) was subsequently conducted to determine the genetic relationships among 147 soybean genotypes via the FactoMineR [28] and FactoExtra R packages [29]. Molecular variance analysis (AMOVA) and calculation of the coefficient of genetic differentiation among populations (PhiPT) were performed to investigate the distribution of genetic diversity among and within hierarchical populations via GenAlEx software (v.6.51) [30].

## Results

### Summary statistics

A total of 59,126 SNP markers from the 147 soybean genotypes were originally generated via the Diversity Arrays Technology (DArT) platform. The transformation of these allelic sequences into genotypic data resulted in a raw data file of 53,418 SNPs, and after quality control analysis (SNP filtering), only 7,083 SNP markers were retained for further analyses. These markers were unequally distributed across the 20 soybean chromosomes (Fig 1; Table 1). The genome-wide SNP density plot indicated that chromosome 18 had the highest concentration of SNPs, accounting for 7.6% of the total number of markers with 538 SNPs. In contrast, chromosome 12 had the lowest concentration, with only 3.17% of the SNPs, totalling 225 markers (Fig 1). The diversity indices for the SNP marker presented a polymorphic information content (PIC) value of 0.277, ranging from 0.262 to 0.293. The MAF averaged 0.254 across all the markers. The observed heterozygosity (Ho) ranged from 0.093 to 0.124, with an average value of 0.110. The expected heterozygosity (He) varied between 0.322 and 0.371, with an average of 0.344 (Table 1, S1 Fig).

**Table 1.**
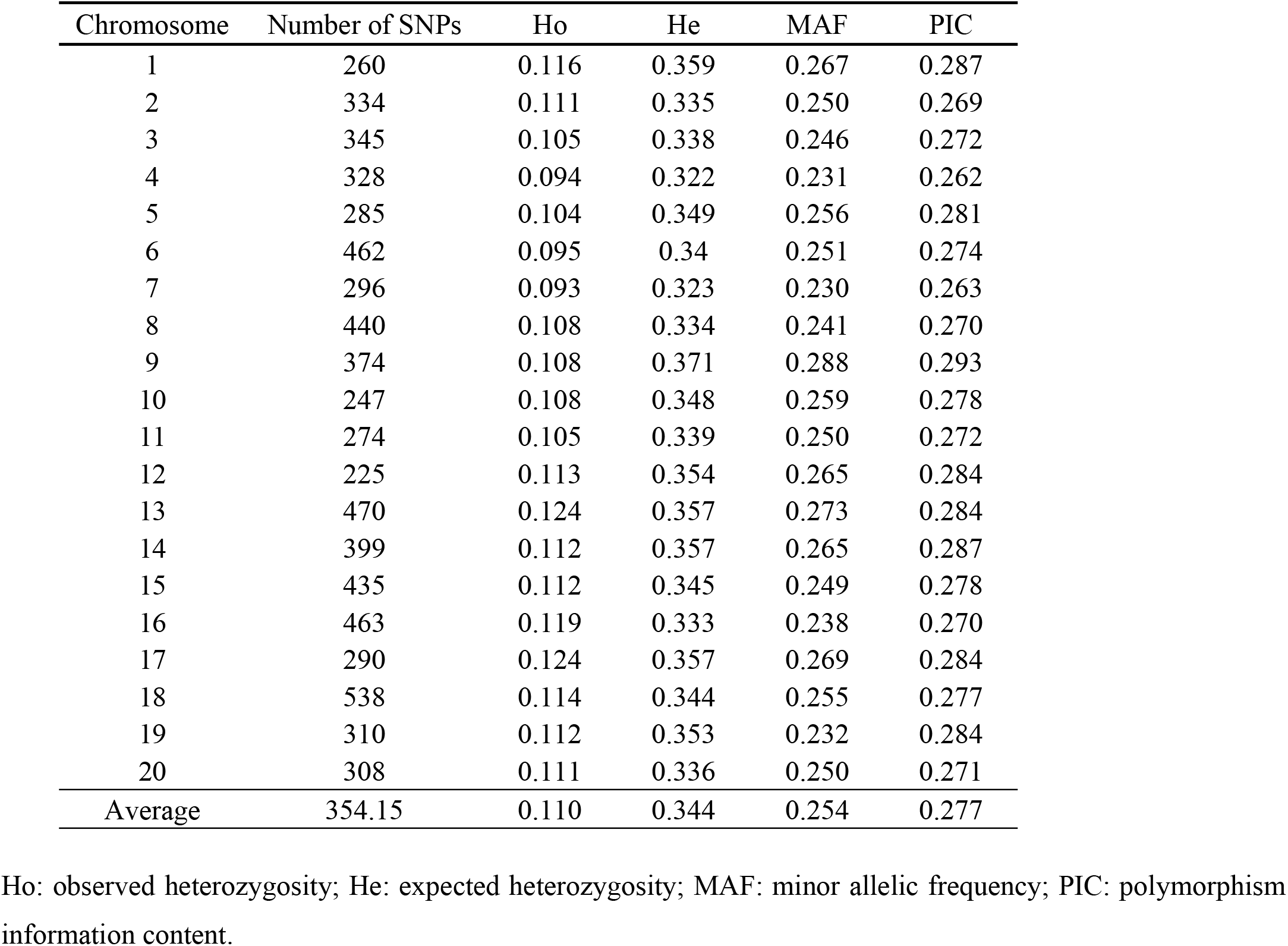
Summary statistics of diversity indices for 147 soybean accessions based on 7,083 SNP markers.

| Chromosome | Number of SNPs | Ho | He | MAF | PIC |
| --- | --- | --- | --- | --- | --- |
| 1 | 260 | 0.116 | 0.359 | 0.267 | 0.287 |
| 2 | 334 | 0.111 | 0.335 | 0.250 | 0.269 |
| 3 | 345 | 0.105 | 0.338 | 0.246 | 0.272 |
| 4 | 328 | 0.094 | 0.322 | 0.231 | 0.262 |
| 5 | 285 | 0.104 | 0.349 | 0.256 | 0.281 |
| 6 | 462 | 0.095 | 0.34 | 0.251 | 0.274 |
| 7 | 296 | 0.093 | 0.323 | 0.230 | 0.263 |
| 8 | 440 | 0.108 | 0.334 | 0.241 | 0.270 |
| 9 | 374 | 0.108 | 0.371 | 0.288 | 0.293 |
| 10 | 247 | 0.108 | 0.348 | 0.259 | 0.278 |
| 11 | 274 | 0.105 | 0.339 | 0.250 | 0.272 |
| 12 | 225 | 0.113 | 0.354 | 0.265 | 0.284 |
| 13 | 470 | 0.124 | 0.357 | 0.273 | 0.284 |
| 14 | 399 | 0.112 | 0.357 | 0.265 | 0.287 |
| 15 | 435 | 0.112 | 0.345 | 0.249 | 0.278 |
| 16 | 463 | 0.119 | 0.333 | 0.238 | 0.270 |
| 17 | 290 | 0.124 | 0.357 | 0.269 | 0.284 |
| 18 | 538 | 0.114 | 0.344 | 0.255 | 0.277 |
| 19 | 310 | 0.112 | 0.353 | 0.232 | 0.284 |
| 20 | 308 | 0.111 | 0.336 | 0.250 | 0.271 |
| Average | 354.15 | 0.110 | 0.344 | 0.254 | 0.277 |

Ho: observed heterozygosity; He: expected heterozygosity; MAF: minor allelic frequency; PIC: polymorphism information content.

**Fig 1.**
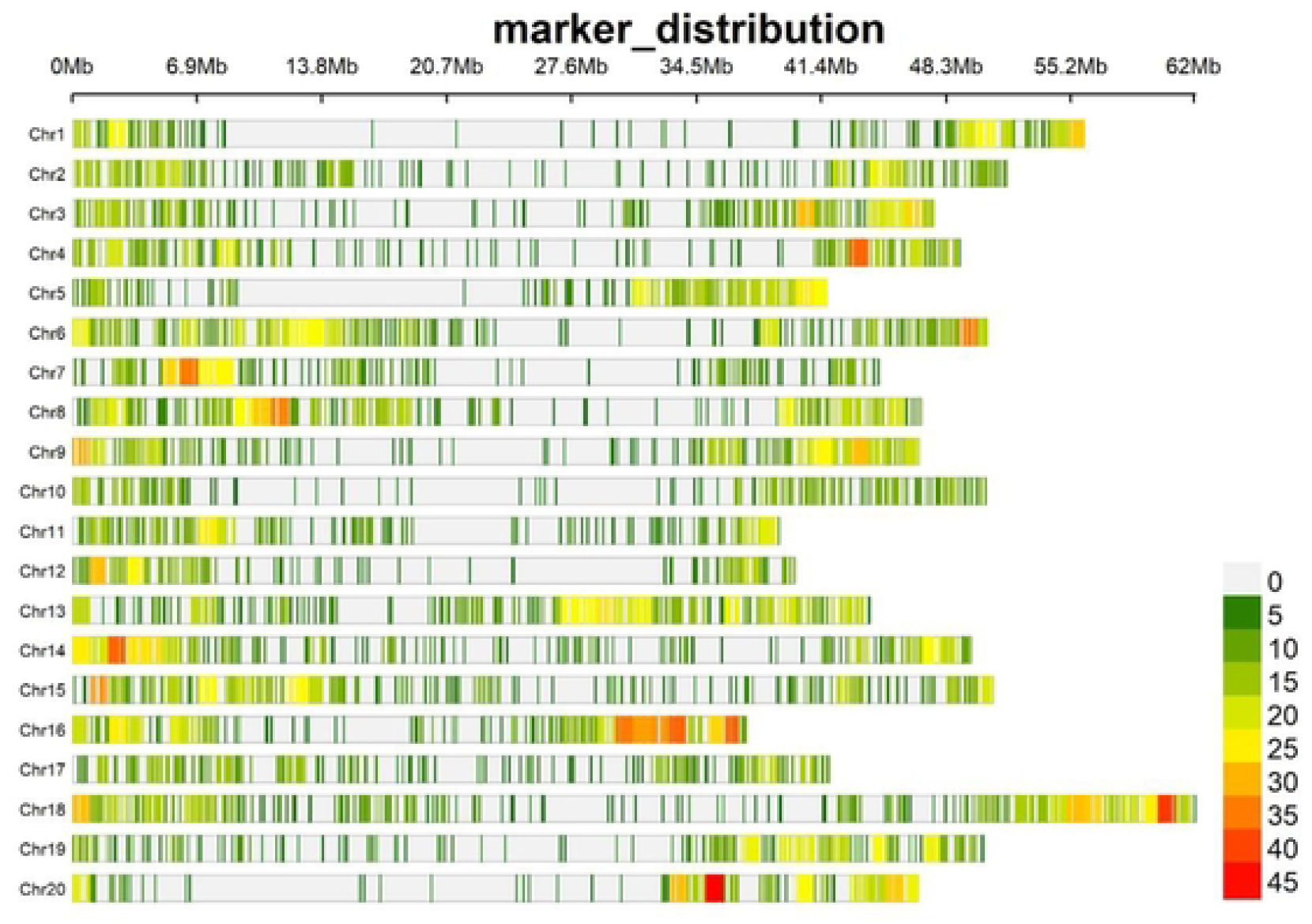
Distribution and density of filtered SNPs across 20 soybean chromosomes. The horizontal axis displays the chromosome length. The number of SNPs density in a given region is indicated at the bottom right.

### Population structure of 147 soybean breeding lines

Various complementary methods, including a model-based Bayesian approach in ADMIXTURE, DAPC, and PCA), were utilized to analyse the population structure of the 147 soybean accessions. On the basis of the admixture results, four subpopulations (K=4) were identified (Fig 2). Similarly, DAPC revealed four genetic groups (Fig 3), following a sharp decline in the Bayesian information criterion (BIC) versus the number of cluster plots (S2 Fig). There was a disparity in how the soybean accessions were assigned to the identified genetic groups between the ADMIXTURE and DAPC results. This disparity may be related to the DAPC analysis, which assigned the 147 soybean genotypes into distinct groups. In contrast, ADMIXTURE assigned 57% of the accessions (84 genotypes) to the four subpopulations on the basis of a membership probability of 70%, whereas the remaining 43% (63 lines) of the collection were classed as admixtures.

**Fig 2.**
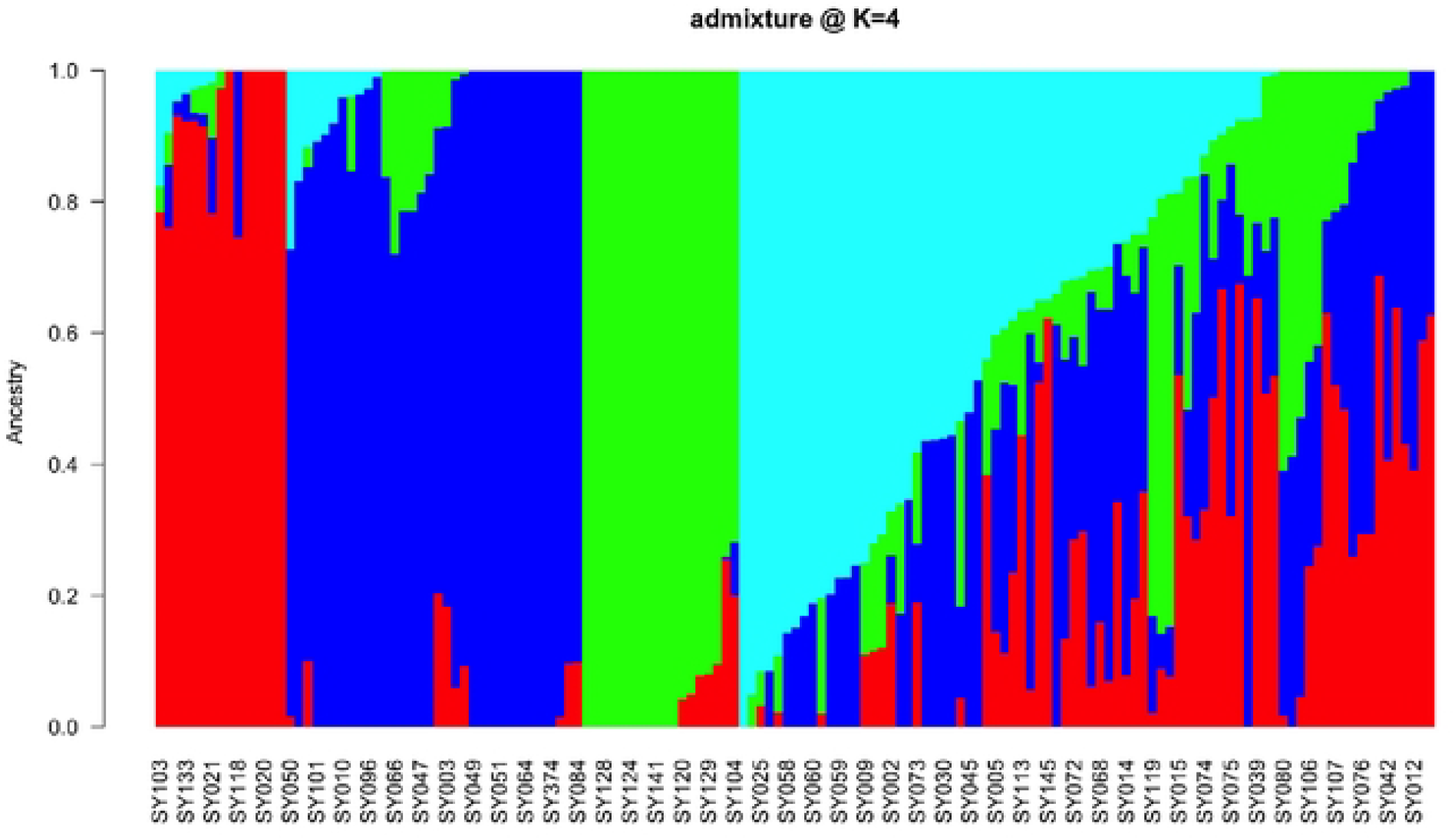
Population structure of 147 soybean breeding lines from the IITA breeding program, Ibadan on the basis of ADMIXTURE analysis with the subpopulations set at K = 4 via 7,083 high-quality SNPs. The colours represent the four subpopulations: Subpopulation 1 (red), subpopulation 2 (blue), subpopulation 3 (green) and subpopulation 4 (cyan), on the basis of a membership coefficient of >70%.

**Fig 3.**
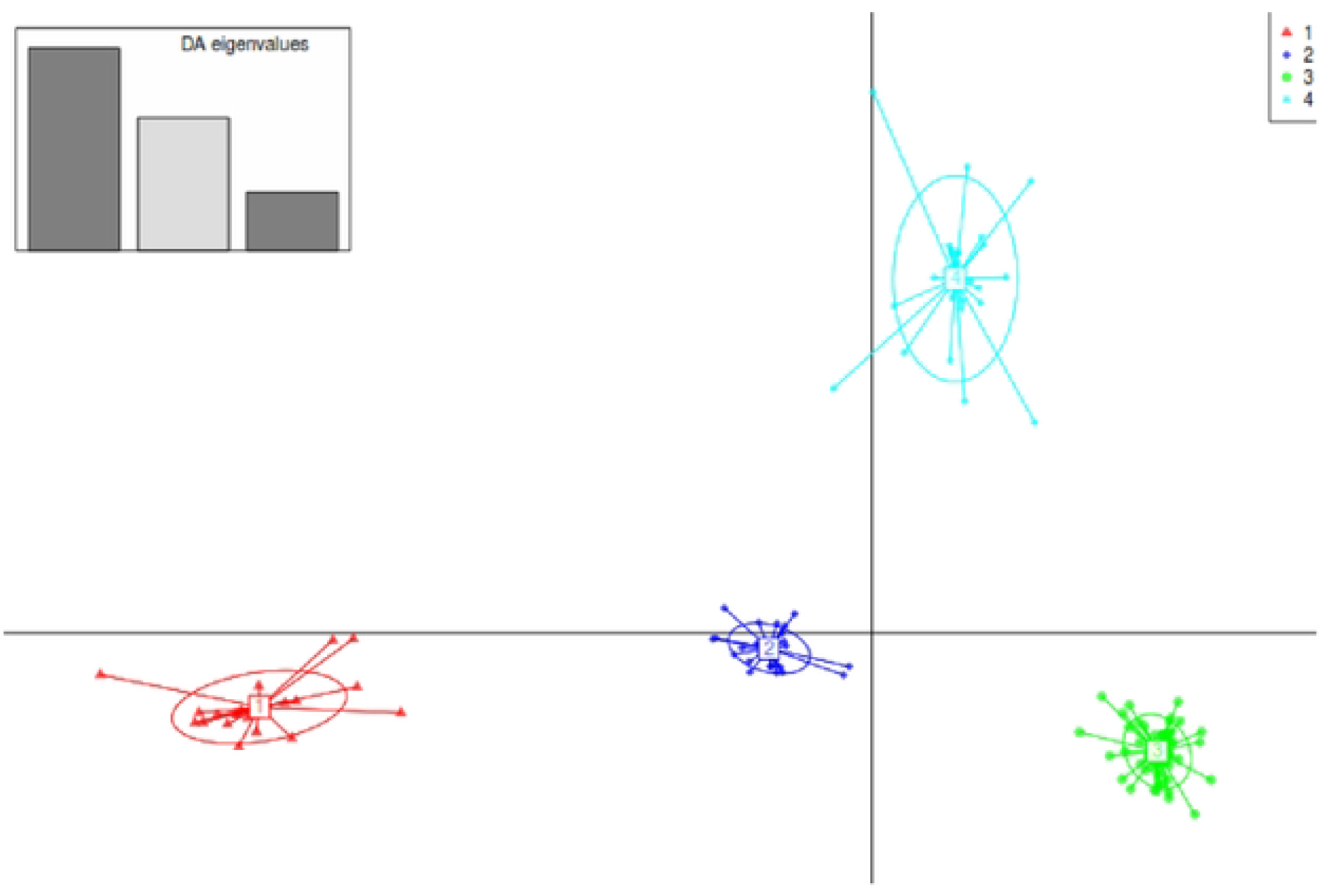
Summary of discriminant analysis of principal component (DAPC) for 147 soybean accessions, illustrating the ordination plot of DAPC for the four groups. Eigenvalues are provided in the upper-left inset. Genetic groups/clusters are represented by different colours and inertia ellipses, with dots indicating individual genotypes.

Hierarchical cluster (HC) analysis grouped all 147 soybean genotypes into three major genetic groups or clusters (Fig 4; S3 Fig). Cluster 1 contained 87 genotypes, predominantly IITA breeding lines, with the exception of a single genotype, ‘SONGDA’ introduced from Ghana, which was originally an IITA breeding line sent to Ghana in variety trials. The 86 IITA genotypes were mainly TGx (Tropical *Glycine max*) varieties or progenies resulting from crosses between two TGx parental lines (Supplementary Table 1). The HC analysis grouped these 87 genotypes into Cluster 1, while the DAPC divided them into two distinct clusters, represented as Clusters 3 and 4 (Figure 3). According to the ADMIXTURE analysis, 36 of the 87 genotypes in Cluster 1, including the unique Ghana genotype, were classified as admixtures. The remaining 51 accessions were assigned to the blue and cyan groups, with 34 and 17 genotypes, respectively (Figure 2). Cluster 2 comprised 34 accessions, including 16 IITA-breeding lines, 16 of the 17 genotypes sourced from the USDA soybean genetic resource center, and one variety Sc-Signa from SeedCo (a private Company) and MAKSOY-4N from Makere University, Uganda. The 16 IITA breeding lines consisted of progenies derived from various parental lines, including TGx, ZIGx, SPSOY, CIMARRONA, PI567090, SOYICA and ST SUPREMA (S1 Table). Among the 34 genotypes in Cluster 2 identified by HC analysis, 15, exclusively IITA breeding lines, were clustered by ADMIXTURE analysis in subpopulation 1 (red) (Figure 2). The remaining 19 genotypes, which included one IITA breeding line, 16 from Columbia, and the unique genotypes from SeedCo and Uganda, were assigned as admixes by ADMIXTURE analysis. On the other hand, the DAPC analysis placed all Cluster 2 genotypes into Cluster 1 (Figure 3). The 26 genotypes assigned to Cluster 3 by HC analysis included 25 IITA breeding lines and one Columbia genotype (Panaroma-3). The IITA breeding lines were a mix of pure TGx parents and backcross progenies, derived from crosses between TGx lines and other parental lines, such as ST SUPREMA, CIMARONA, SOYICA, ZIGx, LG-12, and AS-G (S1 Table). The DAPC analysis classified these 26 genotypes from Cluster 3 in the HC analysis into Cluster 2 (Fig 3). Moreover, the ADMIXTURE analysis placed 18 of them into a specific group (green) (Fig 3), whereas the remaining 8, including the unique USDA genotype (Panaroma-3), were categorized as admixtures.

**Fig 4.**
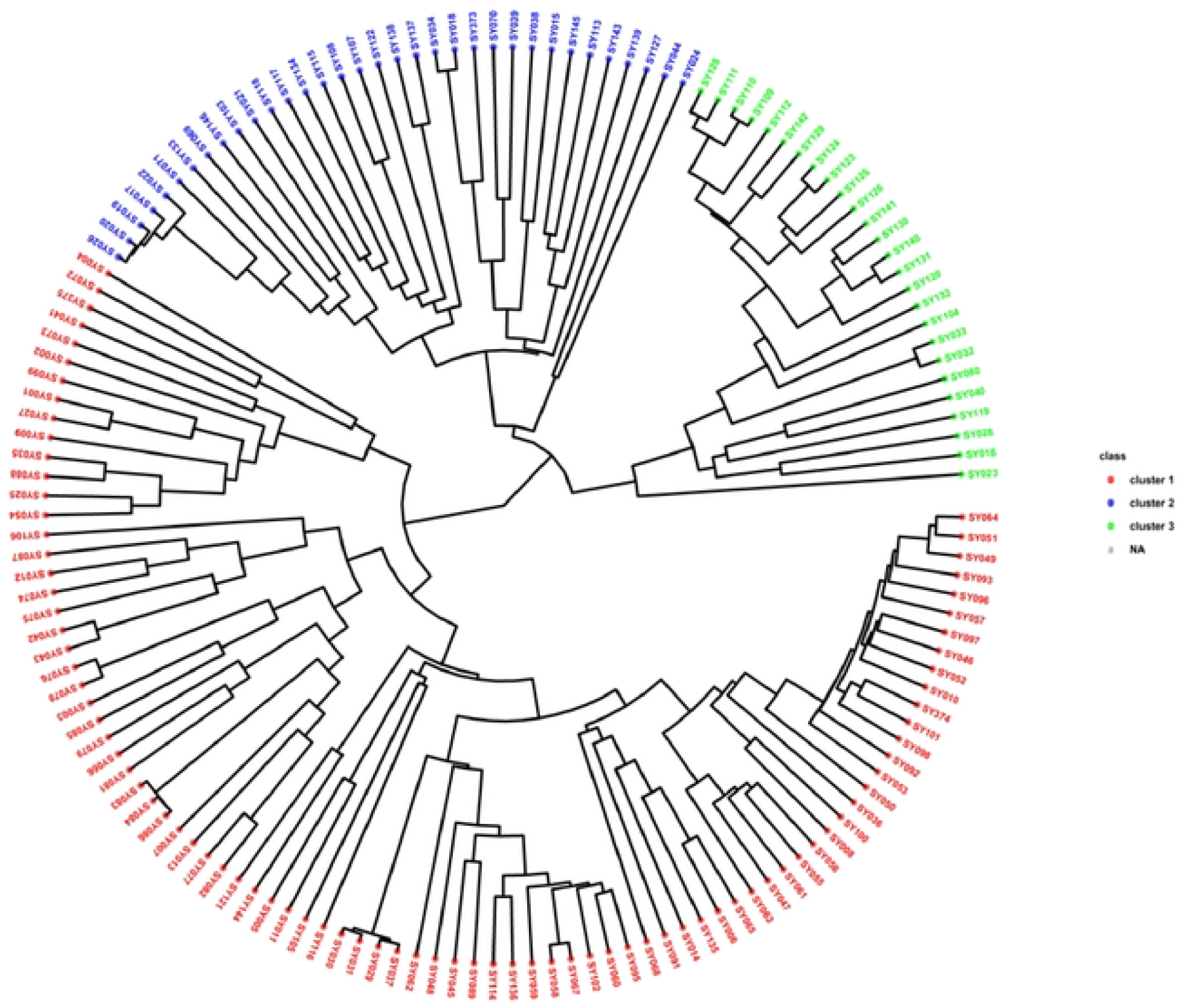
Hierarchical clustering analysis based on 7,083 DArT-SNP markers, depicting the genetic relationships among 147 soybean accessions from the IITA, Ibadan breeding programme.

Principal component analysis (PCA) revealed that the first and second components (PC1 and PC2) accounted for 45.2% and 15.9% of the total molecular variation, respectively, together explaining 61.1% of the overall observed variation (Figure 5). Although all the genotypes within each cluster were grouped, they exhibited some heterogeneity. The genotypes classified as admixtures were identified as admixed groups in the PCA (Figure 5).

**Fig 5.**
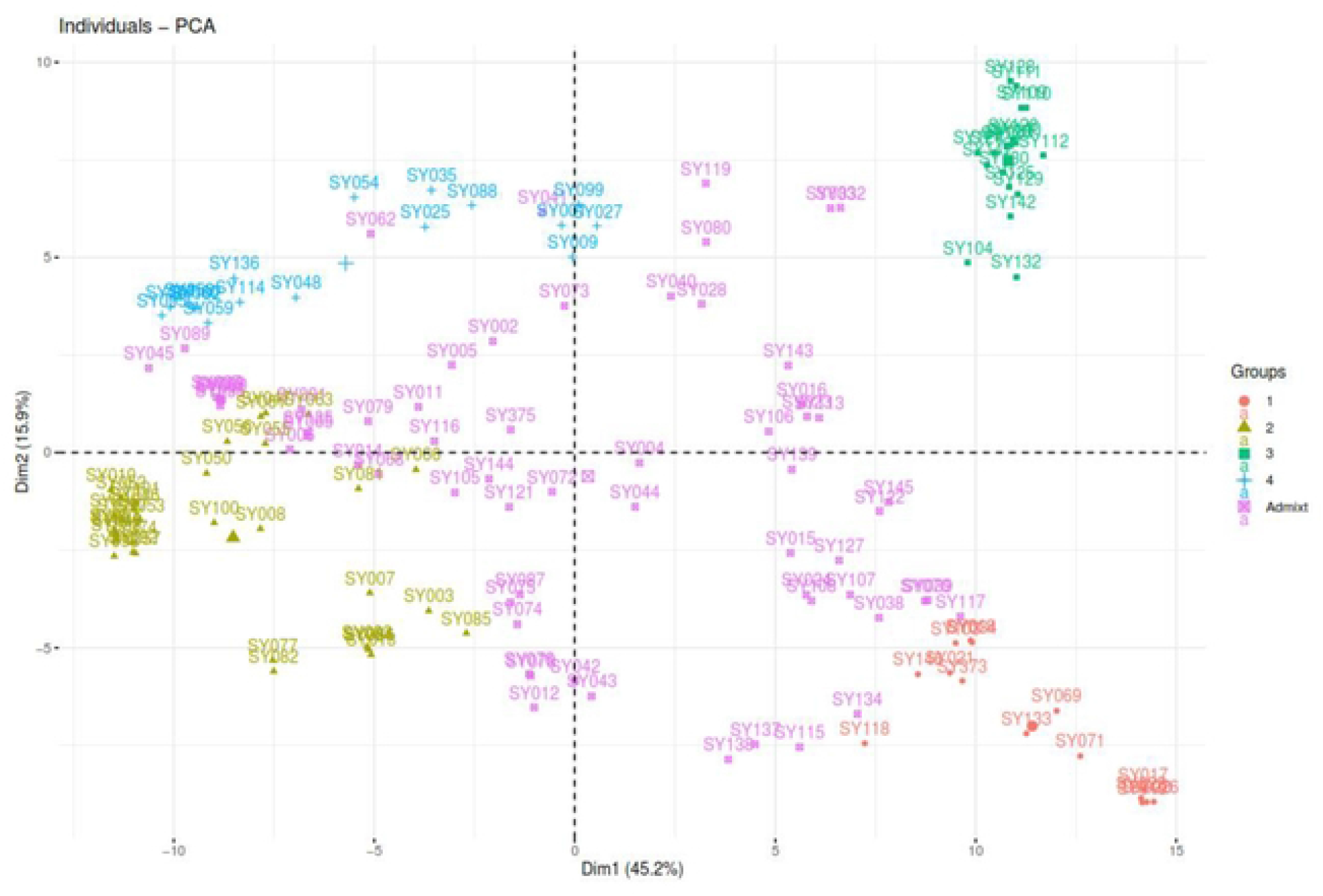
Principal component analysis plot showing the clustering of 147 soybean breeding accessions into four clusters. Each cluster is represented by a color: cluster 1 (red), cluster 2 (yellow), cluster 3 (green) cluster 4 (blue), and admixed (pink).

### Genetic distance and differentiation of soybean accessions

A pairwise dissimilarity genetic distance matrix revealed that the genetic distance among the 147 soybean genotypes ranged from 0.012 to 0.452, with an average distance of 0.333. The greatest genetic distance of 0.452 was found between the USDA genotype TGx 2029-39F (Cluster 2) and two IITA breeding lines, TGx 2002-89 GN and TGx1988-5FxTGx1989-19F-9, both in Cluster 1. In contrast, the lowest genetic distance (0.012) was observed between two IITA lines, TGx 2002-89 GN and TGx 2002-90 GN, both of which belonged to cluster 1. Within Cluster 1, the genetic distances ranged from 0.012 to 0.452 with an average of 0.337. Cluster 2 presented genetic distances ranging from 0.015 to 0.452, with an average of 0.367. For Cluster 3, the distances ranged from 0.017 to 0.433, with an average of 0.355.

The analysis of molecular variance (AMOVA) revealed that 77% of the total genetic variability was partitioned as within-population variation, which was significantly greater than the 23% partitioned among among-populations variation (Table 2). The overall genetic differentiation (PhiPT) and gene flow (Nm) for the 147 soybean genotypes were 0.233 (p < 0.001) and 1.649, respectively.

**Table 2.**
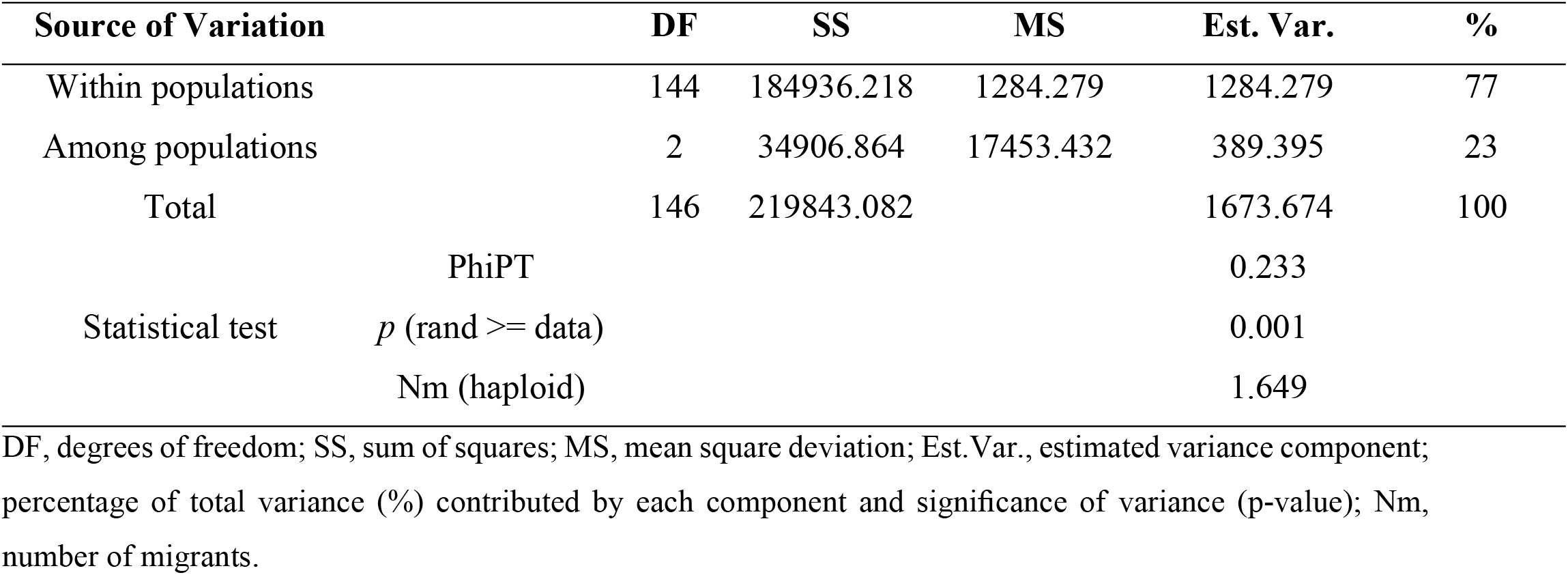
Analysis of molecular variance (AMOVA) within and among soybean populations.

The pairwise population differentiation (PhiPT) estimates revealed that the highest degree of differentiation (0.267) was observed between populations 1 and 3, whereas the lowest degree of differentiation (0.200) occurred between populations 1 and 2. The genetic differentiation between population 2 and population 3 was 0.244. On the other hand, the pairwise population estimates of gene flow (Nm) for the three populations ranged from 1.376 to 1.998 migrants per population (Table 3).

**Table 3:** **Pairwise population PhiPT values and Nm values based on 999 permutations from AMOVA according to HC analysis** (all PhiPT values were significantly greater than 0, p < 0.001).

| Population 1 | Population 2 | PhiPT | Nm |
| --- | --- | --- | --- |
| Cluster 1 | Cluster 2 | 0.200 | 1.998 |
| Cluster 1 | Cluster 3 | 0.267 | 1.376 |
| Cluster 2 | Cluster 3 | 0.244 | 1.546 |

## Discussion

Studying the genetic diversity of germplasm or breeding material is the best approach for understanding the existing genetic variation and effectively managing genetic resources to enhance breeding programs [31, 32]. Hence, plant breeders need such genetic analysis to execute strategic target selection and integration while maintaining significant economic traits associated with distinct crops [33].

The observed PIC ranged from 0.262 to 0.293, with an average value of 0.277, across 7,083 SNP markers on the 20 soybean chromosomes, indicating that these markers are both informative and polymorphic. In a previous study, Botstein et al. (34) reported that SNP markers with PIC values greater than 0.5 are highly informative, those with PIC ranging from 0.25 to 0.5 are informative, and SNPs with PIC values less than 0.25 are fairly informative. The average PIC value found in this study was slightly greater than the reported average values of 0.25, and 0.22 for soybean collection reported by [2] and [35], respectively. This study also demonstrated the possibility of using the selected DArTseq-SNP markers for genomic investigations in soybeans, which may serve as a foundation for future breeding efforts in the IITA soybean breeding program and conservation initiatives in Nigeria. The MAF value measures the selective ability of the marker. Owing to the bi-allelic nature of SNP markers, the MAF closest to 0.5, is best. The high average MAF value of 0.254 observed in this study indicates valuable genes can be exploited from those genotypes [32]. Compared to the results based on SNP markers reported by Hao et al. [36], our MAF values are lower. Their study revealed that the MAF ranged from 0.102 to 0.50 in soybean landraces, with an average value of 0.291. This difference might be because the materials used in the present study were advanced breeding lines, whereas Hao et al. [36] focused mainly on landraces. The average expected heterozygosity (He) of 0.344 indicates high genetic diversity within the soybean accessions, which can be effectively used for soybean improvement [32].

Analysing population structure via SNP markers offers helpful information for preserving and tracking the genetic diversity essential for an effective breeding program [37]. ADMIXTURE and DAPC analyses were used to determine the population structure, revealing the presence of four major populations (K = 4) for the 147 soybean genotypes. However, previous studies [2] and [6] reported different ADMIXTURES results, with ΔK values of 3 and 6, respectively. Considerable levels of admixture (42.87%) were detected among the genotypes, which likely resulted from historical gene flow, breeding practices, and inherent genetic diversity within and between the soybean populations [38]. Chander et al. [6] reported similar levels of admixture in their study of 165 soybean genotypes, which primarily consisted of IITA-bred soybean varieties. In contrast to the results of the ADMIXTURE and DAPC analyses, the hierarchical cluster (HC) method classified the 147 genotypes into three major clusters, which tended to group related lines by their origin or pedigree. These results suggest that the distinct pedigrees of each soybean genotype were crucial in preserving genetic variation, as genotypes with similar pedigrees clustered together on the basis of the SNP markers. Soybean genotypes of diverse origins may present significant genetic differences [39]. The fact that the first two principal components (PCAs) explained 61.1% of the overall genetic variation highlights the effectiveness of highly informative and selective SNP markers for genetic studies in soybeans, which are essential for conservation and future breeding initiatives. Moreover, the PCA demonstrated a stronger association among accessions within each cluster. Our PCA findings indicate that accessions within the same cluster are relatively close and that crosses between accessions from different clusters could lead to increased genetic diversity in IITA’s soybean breeding programs on the basis of the different product profiles.

In this study, AMOVA revealed that 77% of the genetic variance was within populations, whereas 23% was due to variance among populations. Similar findings have been reported in previous studies on soybean [10, 40–42] and other crops, such as *Camelina sativa* [43], wheat [44], rice [14], cowpea [31, 45], and potatoes [46]. The PhiPT value, an analogue of the fixation index F_ST_, indicated high differentiation between populations 1 and 2 and between populations 2 and 3. Additionally, very high differentiation was observed between populations 1 and 3. These significant divergences among all the pairwise populations suggest high diversity among the soybean accessions and highlight the informativeness and utility of the selected markers for future soybean genetic diversity research [2]. Hence, hybridizing genotypes from different populations could introduce the necessary variation to increase genetic gain through active selection [10]. Additionally, gene flow (Nm) enhances the genetic diversity of plant populations and is a crucial factor influencing genetic differentiation [47]. When Nm is greater than 1, gene flow is sufficient to counteract the effects of genetic drift. In the present study, the average gene flow value (Nm = 1.649) observed among the populations indicates that the current soybean populations are not yet impacted by genetic drift.

## Conclusion

In this study, the SNP data obtained through advanced molecular techniques provide valuable insights for soybean breeding and genetic research. The analysis revealed that significant genetic diversity exists among the soybean accessions. This diversity could serve as a foundation for developing new soybean varieties with superior grain yield potential, broad adaptability, and enhanced resistance to both abiotic and biotic stresses. The results also revealed that the accessions could be categorized into four main groups on the basis of the population structure analysis and three groups according to hierarchical cluster analysis. This information on the genetic diversity and population structure of the IITA West Africa soybean gene pool will be applied in both current and future research. This study will aid in the marker-trait discovery of economically valuable traits in soybean for better and faster improvement.

## Supporting information

**S1 Table. List of the soybean (*Glycine max* (L.) Merril) accessions used in this study and the respective origins**

**S1 Fig. Summary statistics of 7**,**083 single nucleotide polymorphism (SNP) markers used for genotyping 147 soybean accessions**. (a) Expected heterozygosity, (b) observed heterozygosity, (c) minor allele frequency and (d) polymorphic information content

**S2 Fig. Graph showing the best k value via Bayesian information criterion analysis. S3 Fig. Silhouette graph showing the optimal number of hierarchical clusters**.

## Acknowledgements

The first author is grateful for the scholarship granted by the African Union Commission for his PhD studies at the Pan African University Life and Earth Sciences Institute (PAULESI), University of Ibadan, Nigeria. We are grateful for the technical support the soybean breeding team provided in establishing the trials and leaf sample collection.

## Author contributions

**Conceptualization**: Tenena Silue, Paterne Angelot Agre, Bunmi Olasanmi, Abush Tesfaye Abebe.

**Data curation**: Tenena Silue.

**Formal analysis**: Tenena Silue, Adeyinka Saburi Adewumi, Paterne Angelot Agre.

**Investigation**: Tenena Silue

**Methodology:** Tenena Silue, Paterne Angelot Agre, Adeyinka Saburi Adewumi, Abush Tesfaye Abebe.

**Resources**: Abush Tesfaye Abebe

**Software**: Tenena Silue, Adeyinka Saburi Adewumi, Paterne Angelot Agre.

**Supervision**: Paterne Angelot Agre, Bunmi Olasanmi, Abush Tesfaye Abebe.

**Validation**: Tenena Silue, Adeyinka Adewumi, Paterne Angelot Agre, Abush Tesfaye Abebe.

**Visualization**: Tenena Silue, Adeyinka Adewumi, Paterne Angelot Agre.

**Writing – original draft**: Tenena Silue

**Writing – review & editing**: Tenena Silue, Adeyinka Saburi Adewumi, Idris Ishola Adejumobi, Paterne Angelot Agre, Bunmi Olasanmi, Abush Tesfaye Abebe.

